# Beyond binding motif: genomic context predicts transcription factor dependent regulation in Arabidopsis

**DOI:** 10.1101/2025.05.23.655699

**Authors:** Laura Turchi, Jérémy Lucas, Gabrielle Tichtinsky, Nicolas Thierry-Mieg, Romain Blanc-Mathieu, François Parcy, Antoine Frénoy

## Abstract

Transcription Factors (TFs) play a crucial role in the spatiotemporal control of gene expression. Despite the presence of many potential TF binding sites (TFBS) across the genome, TFs do not interact with all of them, and, among those, only a subset leads to actual regulation.

In this work, we investigate how the genomic context of a putative TFBS, including the occurrence of binding motifs for other TFs, can be used to predict effective transcriptional regulation by a TF of interest.

We focus on LEAFY (LFY), a plant-specific TF and master regulator of flower development. Using available transcriptomes and TF-DNA binding experiments, we identify 1164 LFY binding sites associated with a regulatory response in *Arabidopsis thaliana*. We then apply a machine learning approach based on properties of the surrounding genomic region, to discriminate these regulatory LFY binding sites from non-regulatory ones. Detailed analysis of the model’s components reveals that, for LFY, the density and quality of binding sites constitute the most important features but were not sufficient on their own to predict regulatory activity. The presence of binding sites for other transcription factors and the overall richness in TFBS were also essential. These results clarify the nature of the regulatory code by which LFY operates and could serve as a basis for the study of the regulatory elements of other transcription factors.

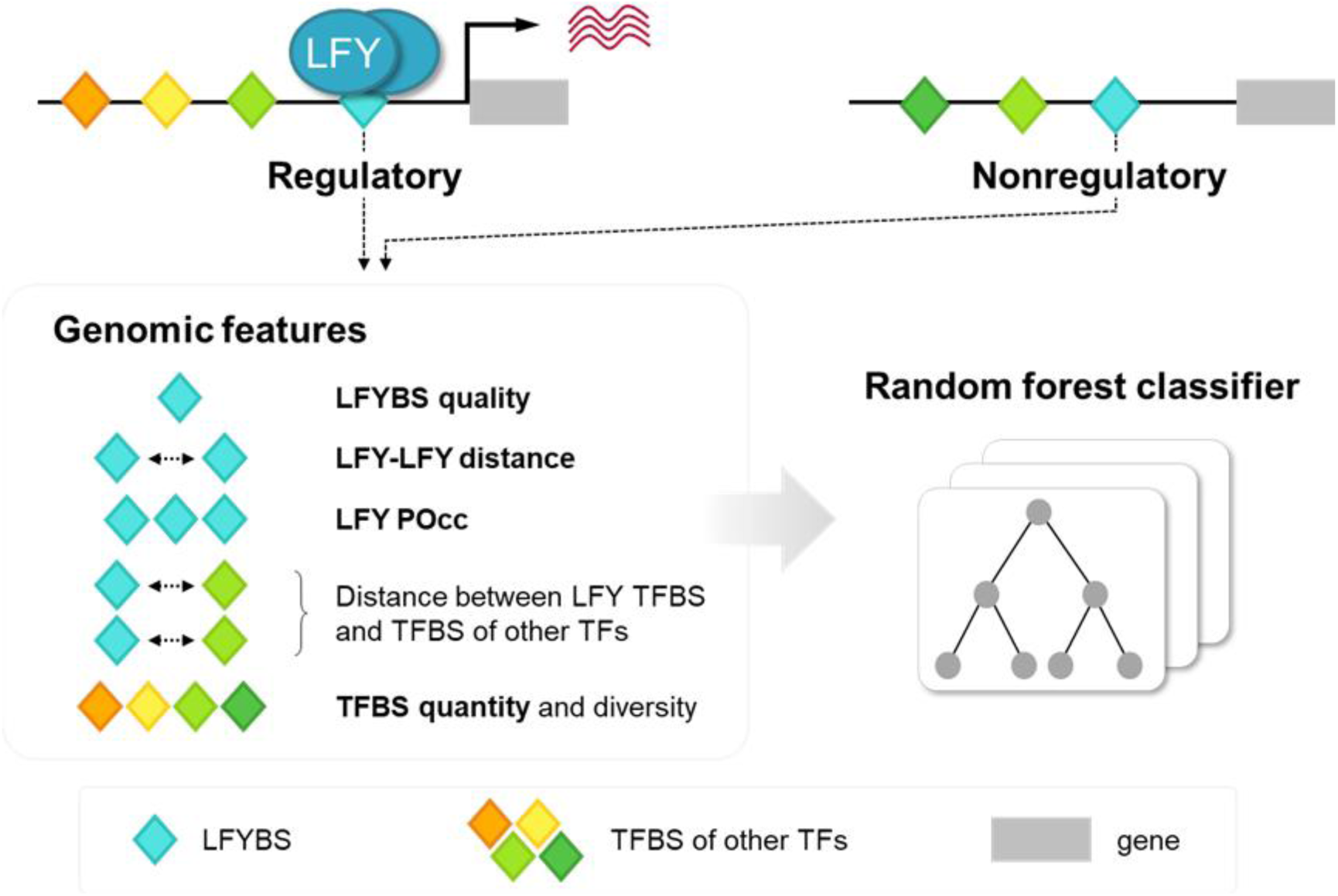

## Introduction

Transcription Factors (TFs) control gene expression by binding to specific DNA sequences (Spitz & Furlong, 2012), and master regulators are TFs that control hundreds of genes to generate important gene expression changes upon developmental transitions (Kaufmann & Airoldi, 2018). Deciphering how TFs recognize their targets and generate a transcriptional response is crucial to understand how they control timely and robust gene expression changes.

A key step of TF-dependent gene regulation is binding to so-called TF binding sites (TFBS) in regulatory regions (Lambert et al., 2018). Experimental approaches such as chromatin immunoprecipitation and sequencing (ChIP-seq) and DNA affinity purification and sequencing (DAP-seq) can provide valuable information about the genome-wide binding profile of a TF *in vivo* and *in vitro*, respectively (O’Malley et al., 2016; Robertson et al., 2007). TF-bound sequences identified by these techniques can be used to determine the specific DNA motif recognized by the protein. An efficient way to represent these DNA motifs is through Position Weight Matrices (PWMs), which contain scores representing how each nucleotide is beneficial or detrimental for TF binding (Stormo, 2015; Wasserman & Sandelin, 2004). The establishment of a large-scale collection of high-quality binding motifs widely adapted by the global community has facilitated genome-wide studies of gene regulation (Rauluseviciute et al., 2024; Sandelin, 2004). PWMs assume that all positions in a binding motif independently contribute to TF-DNA binding, which does not hold true for all TFs (Chen & Pugh, 2021). This has prompted the development of more complex variants of the PWM model such as those including position dependencies (Morgunova & Taipale, 2017; Reiter et al., 2017).

These state-of-the-art DNA binding models nevertheless remain insufficient in predicting TF-dependent regulation (Lai et al., 2019). Multiple TFs can work in a combinatorial fashion to regulate gene expression, and they may require the presence of specific partners in order to fulfill their regulatory role (Morgunova & Taipale, 2017; Reiter et al., 2017; Spitz & Furlong, 2012). This indicates that the genomic context of a binding site contains valuable information such as the potential presence of other TFBSs, which may be important for effective regulation. Yet, individual TFBS motifs cannot capture such genomic context.

The regulatory effect of a TF is typically assessed through the analysis of gene expression changes upon its mutation or overexpression with techniques such as RNA-seq. While these techniques allow direct assessment of effective gene expression, they cannot be used to distinguish direct TF targets from targets of other downstream regulators. Moreover, in multicellular organisms it can be challenging to determine the correct experimental setting, in terms of timing and tissue location, where a TF is expressed and its effects can be detected.

The integration of binding and expression data holds the potential to overcome the limitations of each approach to better characterize the genome-wide targets of a TF.

In this study, we search for the genomic determinants of transcriptional regulation by a TF of interest beyond the mere genomic occurrence of its cognate binding motif. As a case study, we focus on LEAFY (LFY), a plant-specific TF and master regulator of floral development (Moyroud et al., 2010; Rieu et al., 2024; Yamaguchi, 2021).

LFY has been extensively studied through genome-wide binding and expression experiments (Goslin et al., 2017; Jin et al., 2021; Lai et al., 2021; Moyroud et al., 2011; Sayou et al., 2016; Schmid et al., 2003, 2004; William et al., 2004; Winter et al., 2011). Moreover, complex binding models have been specifically adapted to LFY. These include an improvement of the classical PWM to account for dependencies between motif positions (Moyroud et al., 2011), and Predicted Occupancy (POcc), a biophysical model that estimates the occupancy of a TF on a given DNA sequence based on the presence of multiple TF sites, their quality, and the biochemical properties of the protein itself (Minguet et al., 2015; Moyroud et al., 2011).

In addition to the presence and quality of LFY binding sites (LFYBS) predicted by these tools, we were interested in genomic information that could help us predict whether a given site is regulatory. Since transcriptional regulation is highly combinatorial, we hypothesize that the presence of other TFs and their genomic position relative to LFY can contribute to regulation. Therefore, we focus on the distance of other TFBS from LFYBS, their density and diversity, as well as the distance of LFYBS from each other. In addition, we examine the distance of each LFYBS to the closest Transcription Start Site (TSS), which has been shown to be important for some TFs (Rozière et al., 2022; Voichek et al., 2024).

We develop and train a statistical model to distinguish between regulatory and non-regulatory LFYBS among all the putative ones found genome-wide with a PWM. We first integrate public and newly generated binding and expression data to define a reliable set of regulatory and non-regulatory LFYBS in Arabidopsis. With this dataset, we use a supervised machine learning approach (random forest models). Our results indicate that genomic features describing the wider genomic context around each LFYBS allow us to distinguish regulatory LFYBS from non-regulatory ones. Our model shows significant improvements in predicting LFY regulatory sites compared to the sole use of state-of-the-art PWM and POcc models. Feature importance analysis reveals the genomic determinants of TFBS regulation by LFY, which seem to be positively influenced by the presence of other LFYBS nearby, and by the quality of the LFYBS themselves. Interestingly, additional features that we hypothesized would be important for predictions, such as TFBS conservation in the green lineage, did not improve model performance. Taken together, these results indicate that genomic context can be used to discriminate regulatory and non-regulatory LFYBS in Arabidopsis.

## Results

### Defining a set of regulatory LFYBS

We started by determining the position of putative LFYBS genome-wide. We used the best-performing LFY PWM available to scan the Arabidopsis genome for predicted LFYBS (Moyroud et al., 2011). By selecting the top 0.1% of all PWM-calculated LFYBS scores, we identified 119138 putative LFYBS genome-wide (Table 1).

**Table 1:** definition and labeling of the dataset. Numbers indicate the amount of LFYBS labeled as regulatory, nonregulatory or undefined based on their overlap with ChIP-seq and DAP-seq peaks or DEGs positions. LFYBS found in intergenic regions were excluded.

| label | definition | sites (excluding intergenic regions) |
| --- | --- | --- |
| regulatory | $\text{ChIP-seq} \cap \text{DAP-seq} \cap \text{DEGs}$ | 1164 |
| nonregulatory | $\text{Not ChIP-seq} \cap \text{not DAP-seq} \cap \text{not DEGs}$ | 56411 |
| undefined | $\text{ChIP-seq} \cup \text{DAP-seq} \cup \text{DEGs}$ except ‘regulatory’ | 40816 |
|  | tot | 98392 |

Among the sites matching the LFYBS motif, we looked for those playing a regulatory role, i.e. with experimental evidence of binding and expression changes at adjacent genes. First, we excluded the 20746 LFYBS detected in intergenic regions, i.e. LFYBS found over 3 kb upstream of the Transcription Start Site (TSS) and more than 1 kb downstream of the transcription termination site.

We then used public datasets of protein-DNA binding experiments to determine LFY’s binding profile genome-wide. As we were interested in regions bound by a LFY homodimer without any additional partners, we used a LFY DAP-seq experiment to select LFY-only bound regions among ChIP-seq peaks. We established that 25398 of the previously detected putative LFYBS are found within DAP-seq peaks, and 5681 within ChIP-seq peaks (Figure 1). A group of 3990 LFYBS are found both within a ChIP-seq and DAP-seq peak, indicating strong experimental support for *in vivo* and *in vitro* binding of these PWM-identified LFYBS.

**Figure 1:**
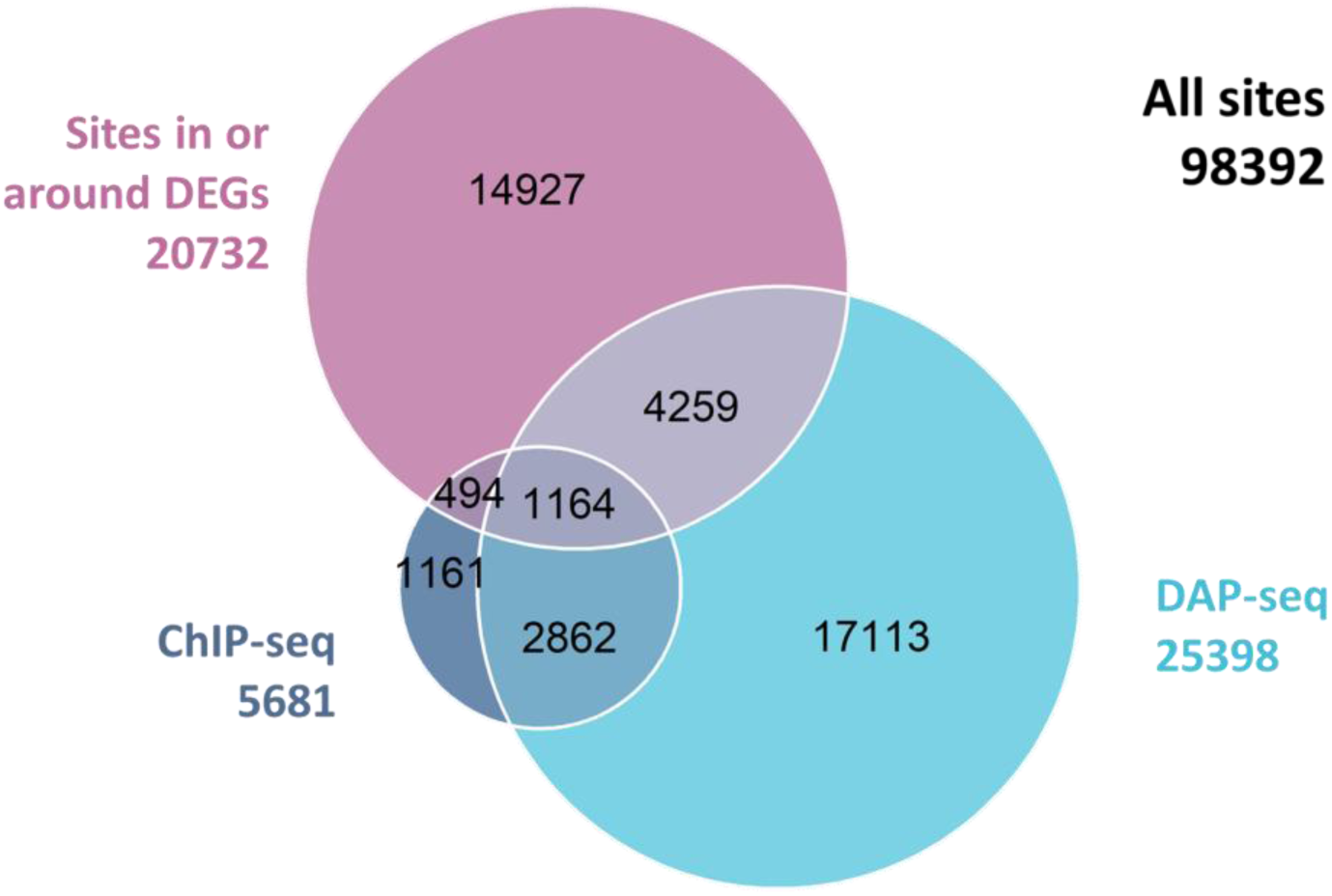
Venn diagram showing the numbers of PWM-detected LFYBS found within ChIP-seq or DAP-seq peaks, or near differentially expressed genes (DEGs). LFYBS close to DEGs were defined as those found within extended DEG regions (extended 3 kb upstream and 1 kb downstream of the annotated gene region).

Binding alone is nonetheless insufficient for effective, *in vivo* transcriptional regulation. We thus integrated the aforementioned binding information with experimental evidence of transcriptional regulation through the analysis of public and newly-generated DNA microarray and RNA-seq experiments. We used experimental settings comparing wild-type plants to *lfy* mutants, experiments where LFY was ectopically and constitutively overexpressed, and LFY inducible systems (Jin et al., 2021; Schmid et al., 2003, 2004; William et al., 2004). We also performed an RNA-seq experiment on wild-type and *lfy* inflorescences (more details in Materials and Methods). By taking the union of differentially expressed genes (DEGs) in these setups, we determined that a total of 20732 putative LFY binding sites are found within or around 3593 DEGs (Figure 1).

Among them, 1164 sites were also bound in ChIP-seq and DAP-seq. We considered these 1164 sites to be high-confidence regulatory LFYBS (Table 1, Figure 1). Conversely, we detected 56411 LFYBS with no evidence of either binding (neither in ChIP-seq nor DAP-seq experiments) or differential expression of nearby genes. We therefore considered them as high confidence nonregulatory LFYBS, i.e. matching the LFY motif but unlikely to contribute to regulation by LFY.

Finally, we found a group of 40816 sites that had partial evidence of regulatory potential, e.g. they were found in regions bound by LFY in ChIP-seq or DAP-seq experiments, or close to DEGs. We classified such sites as ‘undefined’ as we did not have enough evidence to confidently label them as regulatory or nonregulatory (Table 1).

This procedure led us to the establishment of high-confidence positive (regulatory) and negative (nonregulatory) classes of LFYBS.

### Extracting relevant genomic descriptors of genome-wide LFYBS

Using the aforementioned high-confidence labeled datasets, we aimed at finding genomic features predictive of effective transcriptional regulation by LFY, beyond the sole presence of a binding motif. To this end, we computed for each putative site a set of properties which could hold predictive potential.

Firstly, we annotated the type of sequence where each LFYBS is found. We did so by including binary variables describing whether the focal LFYBS lies within a promoter (3 kb region upstream of the TSS), a 5’ untranslated region (UTR), an exon, an intron, a 3’ UTR or downstream (1 kb downstream of the transcription termination site) region. We used six binary variables rather than a single categorical one as a given location can belong to several of these categories.

Then, we hypothesized that the score of the LFYBS could be used as a proxy of LFYBS quality, and thus important in determining its regulatory activity. Therefore, we included the PWM score of each LFYBS. We also included the POcc score in windows of ±250 bp or ±500 bp around each LFYBS, as we thought this alternative measure, which can take into account the existence of several binding sites (and their quality) in the form of a single score value, could capture LFY’s ability to form higher-order oligomeric structures (Sayou et al., 2016). Still in consideration of the cooperative nature of LFY binding, we also included the base-pair distance of each LFYBS from its next closest LFYBS.

Several TFs can co-occur on the same regulatory regions and contribute to regulatory processes in a combinatorial fashion, so we also looked for the presence of TFBS belonging to other TFs around LFYBS. As TFs within the same family tend to have similar binding motifs, we limited redundancy by extracting 46 PWMs representing 46 TF clusters with similar binding motifs from the JASPAR 2022 database (Castro-Mondragon et al., 2022). We scanned the Arabidopsis genome with each one, and we applied the same threshold as LFY (top 0.1%) to determine the presence of a putative TFBS. Then, we computed the distance between each LFY site and the closest site of each cluster.

Regulatory regions are generally enriched in TFBS (Spitz & Furlong, 2012), regardless of the specific identity of these TFs. Thus, we also computed the total amount of non-LFY TFBS around LFYBS (±250 bp or ±500 bp), as it could be an indicator of the presence of a regulatory region and therefore hold predictive potential of regulatory activity.

It has been shown that regulatory regions are not only characterized by a high density of TFBS, but also by TFBS diversity, i.e. the presence of TFBS belonging to diverse TFs (Singh et al., 2021). Therefore, we computed non-LFY TFBS diversity around LFYBS through two different methods. The first was an adaptation of Shannon’s entropy, while the second was a simple count of the amount of non-LFY TFs with at least one TFBS around LFY sites (more details in Materials and Methods). We computed these TFBS diversity scores in ±250 bp or ±500 bp windows around LFYBS.

Moreover, some TF families display TFBS-TSS preferential distances. For instance, the presence of a binding motif of GATA TFs on either side (upstream or downstream) of the TSS has been shown to influence gene expression during Arabidopsis development (Rozière et al., 2022; Voichek et al., 2024). Due to this evidence of the importance of TFBS-TSS distance for gene regulation, we included LFYBS-TSS distance in our model.

Finally, as functionally important TFBS have been reported to be evolutionary conserved along the tree of life, including in plants (Ballester et al., 2014; Berthelot et al., 2018; Velde et al., 2016), we decided to include TFBS conservation as a potential predictor of regulatory activity. We thought that this information would be particularly important for LFY given the remarkable conservation of its protein sequence and binding specificity across the green lineage (Gao et al., 2019; Maizel, 2005; Sayou et al., 2014). Determining the conservation of TFBS, which typically reside in poorly conserved noncoding regions, is a complex task, but there are reports of the conservation of LFYBS in different plant species (Minguet et al., 2015; Moyroud et al., 2011). Therefore, we used public datasets of genome-wide PhastCons and PhyloP conservation scores, calculated on 64 flowering plants (Tian et al., 2020). We computed the level of PhastCons and PhyloP conservation of each LFYBS through three different methods: a simple average of the conservation score found over the LFYBS, and two different weighted average strategies that use the information content of each nucleotide within the LFY binding motif as weight (more details in Materials and Methods).

Overall, we computed a total of 69 features for each LFYBS, summarized in Table 2.

**Table 2:** LFY-specific and genomic context features used in this work. Features in bold are those later selected to train the best-performing model. See Materials and methods and Supp Fig 1 for more details.

| **Feature name** | **Description** |
| --- | --- |
| **Promoter, UTR_5, CDS, intron, UTR_3, downstream** | Binary column describing whether each LFYBS was found within a given type of region, e.g. within an intron. A single LFY site can overlap multiple sequence types.<br>Promoter: 3 kb region upstream of TSS; downstream: 1 kb region downstream of Transcription Termination Site. |
| **pocc_250, pocc_500** | POcc score calculated $\pm 250$ bp or $\pm 500$ bp around each LFYBS, as in (Moyroud et al., 2011). POcc accounts for the presence of multiple sites within a DNA sequence and for the biochemical properties of the LFY protein. |
| **PWM** | PWM score associated with the LFYBS. Higher scores denote LFY sites of higher quality, with higher probability of binding. |
| **LFY_dist** | Distance between each LFYBS and its closest LFYBS (in bp). |
| **cluster_1-46** | Distance between each LFY site and the closest site of each cluster (in bp). Clusters were taken from (Castro-Mondragon et al., 2022) and represent TF groups of variable size, containing from 1 to 58 different TFs. A total of 46 clusters were used in this study, as one of the 47 clusters computed by Castro-Mondragon et al. only contained LFY and LFYBS-LFYBS distance was computed separately. |
| **TSS** | Distance between each LFY site and the closest TSS (in bp). |
| **TFBS_tot_250, TFBS_tot_500** | Total number of non-LFY TFBS found $\pm 250$ bp or $\pm 500$ bp around each LFYBS. Non-LFY TFBS were defined as TFBS of TF clusters. |
| **Sindex_250, Sindex_500** | Value of Shannon's entropy calculated $\pm 250$ bp or $\pm 500$ bp around each LEAFY site. We adapted the equation to compute Shannon's entropy (Shannon, 1948) to determine non-LFY TFBS diversity around LFY sites (more details in Materials and Methods). |
| **diff_TFs_250, diff_TFs_500** | Total number of diverse non-LFY TFs with a TFBS found within $\pm 250$ bp or $\pm 500$ bp around each LFYBS. Unlike for the 'total number of non-LFY TFBS' feature, this feature quantifies TFBS diversity so multiple TFBS of the same TF were considered as one. |
| **PhastCons, PhyloP** | Average or weighted average of PhastCons and PhyloP conservation scores calculated on each LFYBS. Genome-wide scores were taken from (Tian et al., 2020). More details in Materials and Methods. |

### Training a supervised model to predict regulation by LFY

Current knowledge regarding the combinatorial nature of transcriptional regulation suggests non-linear interactions between different TFs, which implies that purely additive linear models would perform poorly. On the other hand, our data (68 features for 57575 observations, only 1164 of which are positive) does not match the requirements of deep learning. We hypothesized that random forests would represent a reasonable middle ground, especially considering that they can handle datasets that contain both binary and quantitative features and that they allow some level of model explicability through established methods (Breiman, 2001).

We split our dataset into a training set containing 80% of our data, and a validation set with the remaining 20%.

Among the features mentioned in Table 1, four are present in several alternative versions or depend on a parameter which needs to be chosen: window size to compute the total number of TFBS (250 or 500 bp), window size to compute Shannon’s entropy and TFBS diversity (250 or 500 bp), window size to compute Pocc (250 or 500 bp), and measure of conservation (average or weighted average conservation score, or no conservation whatsoever).

In addition to using the training set to define the best model hyperparameters for our data as classically done, we also used it to determine the best version of each of these four features: this choice of version is thus effectively treated as a hyperparameter of the model.

Following this rationale, we used a stratified cross-validation strategy to optimize model hyperparameters (Supp. Table 1) and the choice of alternative feature versions on our training set (Supp. Table 2). The combination of feature versions and hyperparameters which yield the best performance are shown on Supp Fig 1. Additionally, chosen features are represented in bold in Table 2, and are further discussed in the next section of this article. Interestingly, excluding all conservation features improved predictions, thus the final model does not include TFBS conservation.

Using this set of features and hyperparameters, we trained a single model on our entire training dataset, and we tested it on the holdout 20% validation set.

To determine whether our model was capable of predicting if unseen LFYBS were regulatory or not, we constructed a Precision-Recall (PR) curve on the validation dataset and we computed average precision as an approximation of the area under the PR curve. This decision was dictated by the fact that our data is strongly unbalanced, with very few positives against a large number of negatives, which could lead to an overinflation of the commonly reported Area Under the Receiver Operating Characteristic curve (AUROC) (Saito & Rehmsmeier, 2015).

Our model outperforms by far the control model trained exclusively on state-of-the-art LFY PWM and POcc features (Figure 2). This indicates that it is indeed capable of using genomic context information (through the features described above) to predict whether a LFYBS is regulatory or not.

**Figure 2:**
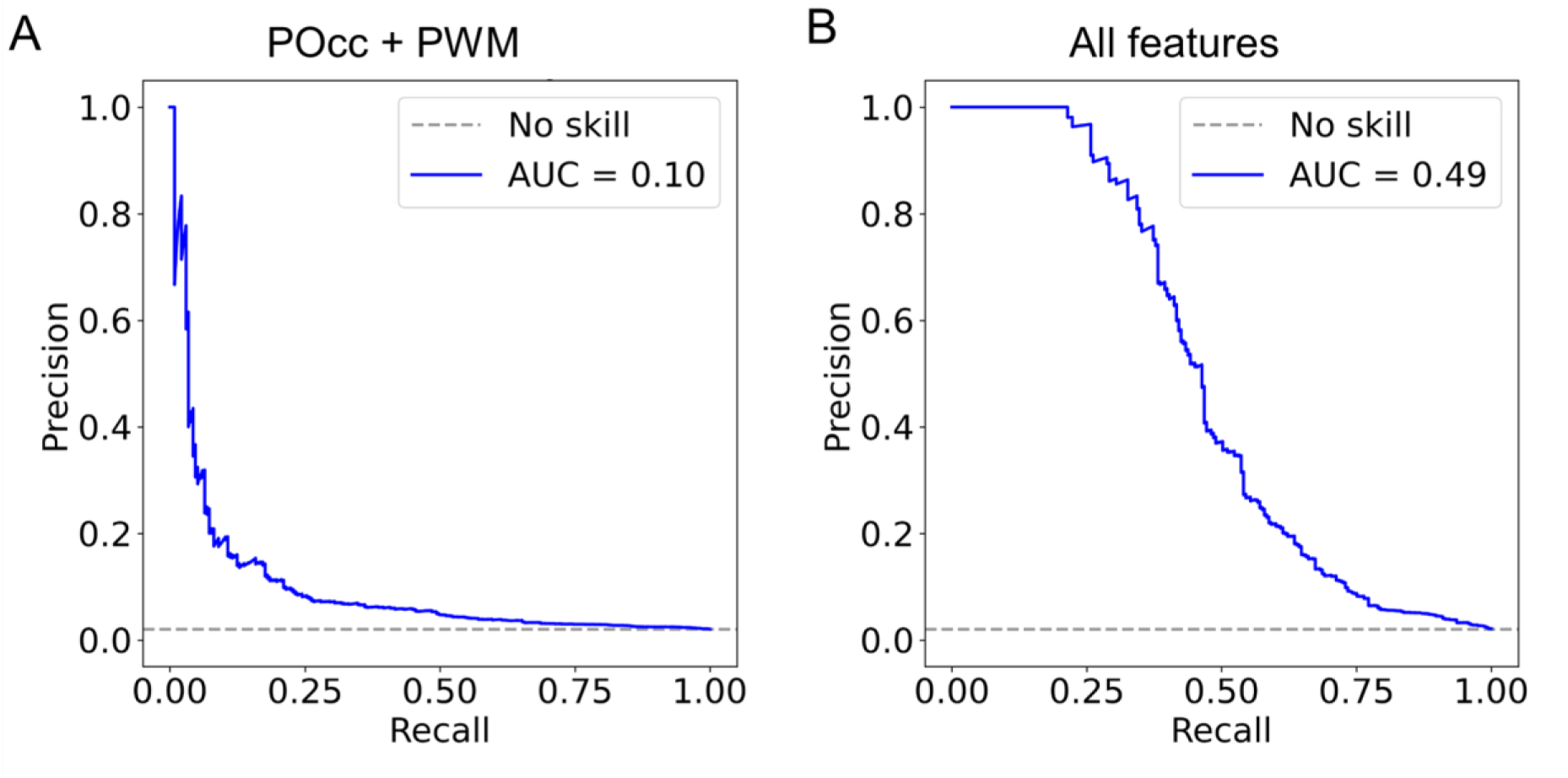
Precision-Recall curve obtained from testing our model on a held-out dataset (20% of our data) after training it on 80% of our data. A: Performance of a model that exclusively uses POcc and PWM features. B: Performance of our model with features highlighted in bold in Table 2. Dashed gray lines represent the performance of a random (‘No skill’) model.

In addition to random forest, we also tested logistic regression and support vector machine (SVM) models on our data, and found that random forest consistently outperformed these models (Supp Fig 1). The superiority of the random forest model could be due to its ability to better capture non-linear relationships and handle interactions between features without the need for explicit feature engineering.

### Analysis of the model

Our results indicate that the best model contains a total of 58 features.

Once we established that our model performed well on the validation dataset, we wanted to find out which features are most important to predict the label of LFYBS. Ultimately, we reasoned that such genomic features are likely to contribute to the regulatory function of LFY in Arabidopsis, thus helping us answer our initial question regarding the genomic determinants of transcriptional regulation.

Therefore, we implemented a permutation-based strategy to determine which features carry important information, which is among the best practices for determining feature importance in random forests (Breiman, 2001). It relies on the permutation of one feature column, and it is quantified as the difference in performance score (in our case, average precision) before and after permutation. Permutation is performed multiple times and for all features.

Based on this strategy, our results show that the three most important features to predict whether a LFYBS is regulatory are all related to LFY: POcc score computed ±250 bp around each LFYBS, distance from the closest next LFYBS, and quality of the LFYBS itself in the form of PWM score (Figure 3).

**Figure 3:**
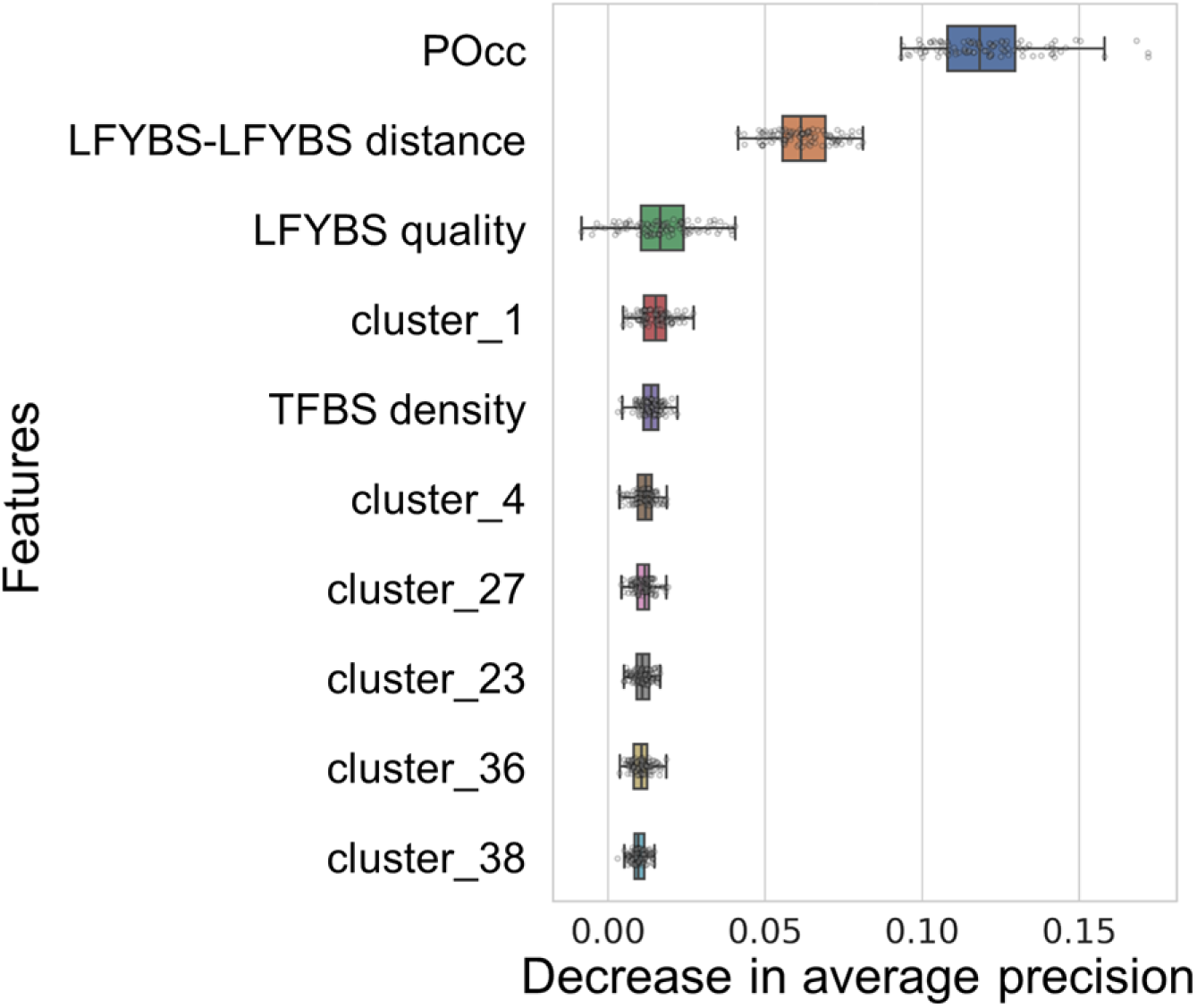
Ten most important features included in our random forest model, as quantified through the decrease in average precision score with a permutation strategy.

While LFY-related features were by far the most important ones, they nevertheless hold little predictive power by themselves (Supp Fig 3). Other important features include the distance of LFYBS from specific TF clusters (namely clusters 1, 4, 27, 23, 36 and 38, Table 3) and the number of non-LFY TFBS present around each LFYBS (Figure 3, Table 3). Each of these features has relatively small permutation importance by itself, but together they bring significant predictive power.

**Table 3:**
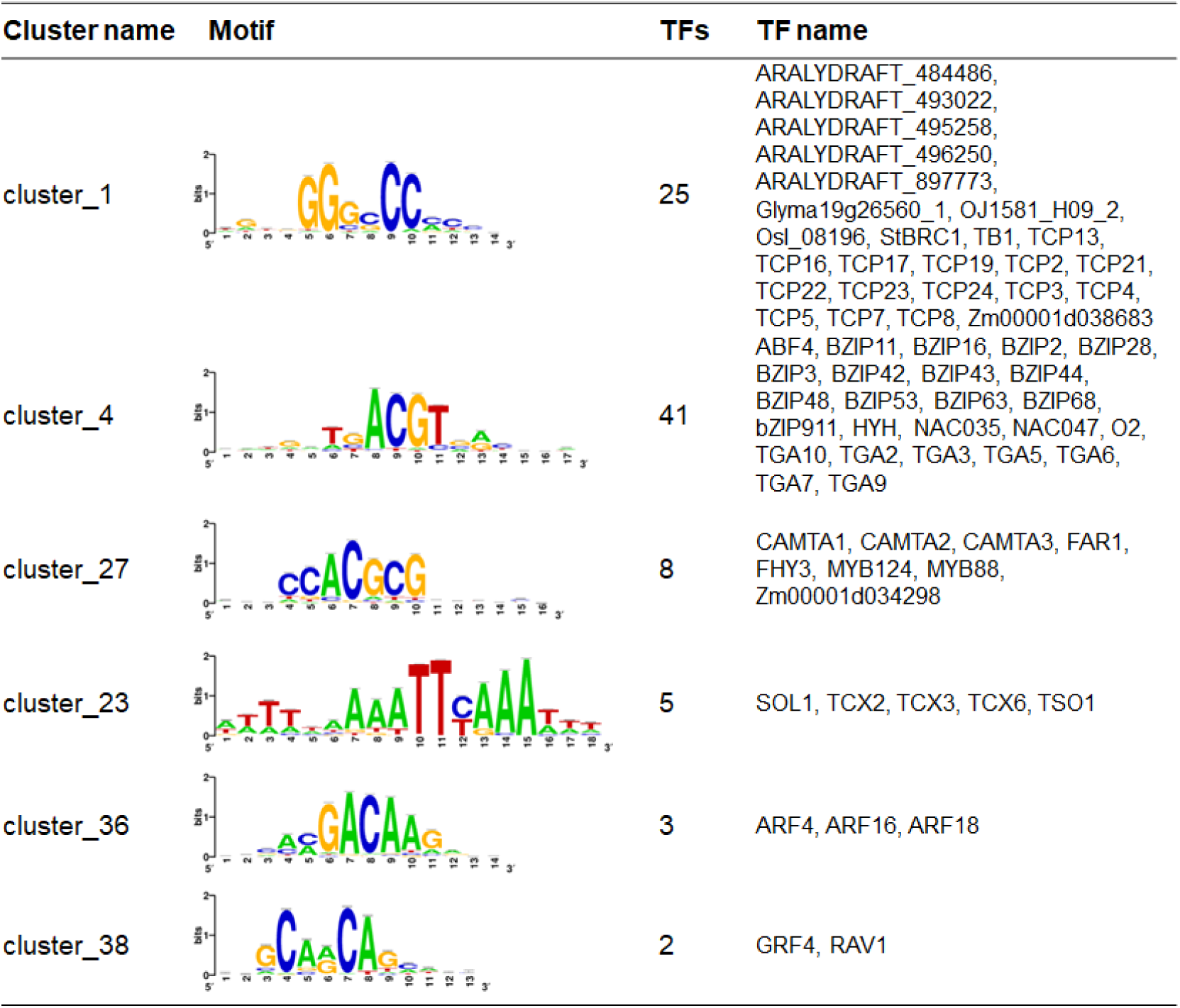
TF clusters in top 10 most important features (Figure 3) from JASPAR 2022 plant CORE collection (Castro-Mondragon et al., 2022). Columns indicate cluster name, its associated motif, total number of TFs in the cluster and their names

Other classical measures of feature importance exist, notably the Gini importance score (Breiman, 2001). We thus also computed Gini scores as a complementary measure of feature importance to the permutation strategy described above. Many of the features with a high permutation score were also found among the highest-scoring features based on Gini importance scores (Supp Fig 3), and although they were not found in the exact same order, they were still correlated overall. This is consistent with previous reports of correlation between permutation- and Gini-based feature importance measures (Strobl et al., 2008).

Other features such as sequence type tended to score very low in our importance scores (Supp Fig 4, Supp Fig 5). This suggests that, unlike more granular information about TFBS content, genomic location is not extremely informative *per se*.

## Discussion

Our results show that, to predict effective regulation by LFY, a model based on genomic context clearly outperforms a model that only uses information related to the presence of a LFY binding motif (PWM or POcc). This confirms our starting hypothesis that binding alone is a poor predictor of transcriptional regulation, and that the highly combinatorial nature of this regulation makes it dependent on the broader genomic context.

Analysis of feature importance indicates that the density and quality of LFY binding sites are the most important for predictions, although they are not sufficient to achieve high model performance (Figure 2, 3). This is consistent with the ability of LFY dimers to form higher-order structures through their SAM domain, which could strengthen binding thanks to clustered LFYBS (Sayou et al., 2016). Groups of TFBS belonging to the same TF are commonly found in animal cis-regulatory regions, where they have been successfully used to elucidate the mechanisms of gene regulation (Gotea et al., 2010). Cooperative binding is not a unique feature of LFY, but rather one shared by many other Arabidopsis TFs. These include TF families with important roles in plant development, such as AUXIN RESPONSIVE FACTORS (ARFs) and MADS box TFs (Freire-Rios et al., 2020; Jack, 2001). Our model was built specifically for LFY, but the same strategy could be applied to other TFs. This would reveal whether this is a particular characteristic of LFY or a common occurrence among other cooperative binding TFs, or even TFs with no evidence of cooperative binding.

In addition to LFY-related features, the distance between LFYBS and the sites of TFs belonging to the plant-specific Teosinte-branched 1/Cycloidea/Proliferating (TCP) family, were also important for predictions (Figure 3). TCP TFs have been shown to be involved in many processes including plant development and flowering (D. Li et al., 2019; S. Li, 2015).

Other groups of TFs also appear among the most important features (Figure 3), including clusters 4, 27, 23, 36 and 38. While no particular group of TFs is dramatically influential on its own, many have a small influence on predictions, suggesting that different combinations of co-occurring TFBS could be contributing to LFY’s regulatory role. The total number of TFBS around LFY sites was also among the most important features for predictions (Figure 3), which suggests that regulatory LFYBS are found in TFBS-rich regions.

Interestingly, sequence type features are among the lowest-ranking ones in terms of importance (Supp. Fig 4 and 5). This suggests that regulatory importance is not always determined by genomic positioning itself, but rather by the regulatory elements it harbors.

Surprisingly, in our setup, the inclusion of conservation features consistently led to lower performance (Supp Fig 2). This goes against widespread reports of the conservation of important regulatory regions (Ballester et al., 2014; Berthelot et al., 2018; Burgess & Freeling, 2014; Velde et al., 2014), under the general rationale that evolutionary conservation indicates purifying selection. There could be several explanations for our contradictory observation. One possibility is that the position of LFYBS differs in more distant plant species. As a consequence, the complex alignment strategy used to compute conservation scores may have failed to capture such sites in divergent, often poorly conserved, noncoding sequences. This would be consistent with reports of LFY-dependent *APETALA1* regulation relying on TFBS that are typically found in promoter sequences in Brassicaceae, while in the more distant Brassicales species *Carica papaya* a putatively regulatory LFYBS is found within the coding region (Minguet et al., 2015). Moreover, LFY regulation has been suggested to be overall conserved despite changes in the position of single LFYBS in regulatory regions (Moyroud et al., 2011).

We chose to only focus on LFYBS that we could assign as regulatory or nonregulatory with solid experimental evidence. As a result, we excluded from our model an important fraction of LFYBS, which we called ‘undefined’ due to incomplete evidence of LFY regulation. Such LFYBS display only binding, *in vitro* or *in vivo*, or evidence of gene expression changes. It is likely that at least part of the ‘undefined’ LFYBS are actually regulatory, but we are lacking evidence. This could be due to technical limitations associated with the nature of the binding and expression data we used. In particular, the effect of *lfy* mutation may be only visible in specific tissues at a specific time, and gene expression datasets are thus likely to contain false negatives. Our trained model could be used to make predictions on ‘undefined’ LFYBS, and predictions would have to be experimentally validated.

Overall, our study provides a successful approach to study the regulatory function of TFs with a supervised model that relies on the presence and location of TFBS on the genome. Besides, our results show that, in addition to being used to predict the regulatory function of TFBS genome-wide, machine learning models can in turn be used to answer fundamental questions about TF functioning in vivo. The main drawback of our approach is that it ultimately relies on PWMs, which have been shown to have limitations (Mathelier & Wasserman, 2013; Siddharthan, 2010; Tognon et al., 2023), and that it requires a significant amount of data, which is not available for all TFs.

On the other hand, the strength of our approach is that it can be flexibly adapted based on the underlying biological question. In our case, as we were interested in finding the genomic properties of LFY regulation, we focused on LFYBS bound both *in vitro* and *in vivo*. This allowed us to exclude targets that require the presence of its known cofactor UFO (Rieu et al., 2023). By doing so, we also excluded targets where LFY binds with other unknown interactors. We propose that a similar approach could be used e.g. to identify new putative interactors by changing the labeling strategy of the input dataset. It would be possible to focus on regions regulated by a protein of interest but without focusing specifically on regions also bound by the protein *in vitro*, as we did by including DAP-seq data. Moreover, our approach requires different types of binding and expression data that are not available for all proteins. When data availability is limited, our approach could be adapted by e.g. focusing only on binding data, while still providing valuable biological insights. Yet again, our work on LFY could also be applied to other TFs, or species that are not Arabidopsis. Additional studies are required to investigate the potential of our approach for predicting regulatory sites when binding and expression experimental data is not available, e.g. in nonmodel species from the genomic context using a model pre-trained on Arabidopsis.

## Material and methods

### ChIP-seq and DAP-seq analysis

LFY ChIP-seq data from (Goslin et al., 2017; Jin et al., 2021; Moyroud et al., 2011; Sayou et al., 2016) were downloaded from GEO (GSE96806, GSE141706, GSE24568, GSE64245, respectively). LFY DAP-seq data was from GEO GSE160013 (Lai et al., 2021). Fastq files were processed as in (Rieu et al., 2023). Briefly, sequencing data quality was evaluated with fastQC v.0.11.7, and adapters were removed with NGmerge v.0.2_dev (Gaspar, 2018). Bowtie2 v.2.3.4.1 was used to map reads to the TAIR10 *A. thaliana* reference genome (Lamesch et al., 2012). We only retained reads mapped to a single location and with a maximum of two mismatches. We used samtools dedup v.1.8 to remove duplicates. We identified bound regions (“peaks”) with MACS2 v.2.2.7.1 (Zhang et al., 2008), with input DNA from Lai et al. as a control (Lai et al., 2021) and with –q 0.05 for ChIP-seq and -q 0.0001 for DAP-seq. Consensus peaks called in all replicates were identified with MSPC v.4.0.0 (Jalili et al., 2015) and, finally, peaks were resized around the peak maximum (±200 bp) for further analysis.

### Microarrays and RNA-seq analysis

Microarray data were retrieved from GEO (GSE28062) for *35S::LFY-GR* seedlings after the addition of dexamethasone (Winter et al., 2011), from (Chahtane et al., 2013) for *35S::LFY* experiments and from AtGen-Express (Schmid et al., 2004) for inflorescence tissue in the *lfy* background. Each mutant or overexpressing genotype was compared to WT (Col-0). The R package gcrma (Wu & Irizarry, 2022) was used to adjust probe intensities and convert them to expression measures, and then the ‘limma’ package (Ritchie et al., 2015) was used for differential expression analysis. Benjamini–Hochberg correction was applied to the P values, and fold change (FC) was computed as the ratio between expression in the overexpressing line/WT (for *35S::LFY-GR* and *35S::LFY*) or WT/*lfy* mutant. Only genes with |log2(FC)| > 1 and adjusted P < 0.05 were considered as significantly differentially expressed.

We performed a new RNA-seq experiment using WT (Col-0) and *lfy-12* plants grown for 5 weeks in short day (SD) conditions (8 h light, 16 h dark), then moved to long days (LD) conditions (16 h light, 8 h dark) for another two weeks before inflorescence dissection. All flower-looking structures were removed, and only the upper part of the inflorescence was sampled (2-3 mm). We only sampled primary inflorescences for a total of four samples per genotype, each sample containing multiple inflorescences. RNA was extracted with the Qiagen Rneasy Kit and sent for paired-end mRNA sequencing.

Our newly produced RNA-seq samples, like RNA-seq data of dexamethasone-treated *35S::LFY-GR* calli published in (Jin et al., 2021), were analyzed as follows. Fastq files were trimmed with BBduk to remove adapter sequences. Reads were mapped to the TAIR10 genome with STAR (Dobin et al., 2013). The tximport R package (Soneson et al., 2015) was used to import read counts to RStudio v1.3.959 (RStudio Team, 2020), and the DESeq2 package v1.28.1 (Love et al., 2014) was used to run differential expression analysis. Genes with |log2(FC)| > 1 and adjusted P < 0.05 were considered as significantly differentially expressed.

### Identifying LFYBS genome-wide

A LFY PWM with dependencies (Moyroud et al., 2011) was used to scan the *A. thaliana* genome (TAIR10) (Lamesch et al., 2012) to predict LFYBS, and a PWM score was assigned to every genomic position. All genomic scores were used to determine an overall distribution, which was then used to select LFYBS based on percentile thresholds. We only retained the top 0.1% best-scoring LFYBS genome-wide.

### Labeling regulatory/nonregulatory/‘undefined’ LFYBS

We integrated binding (ChIP-seq and DAP-seq) and expression (microarrays and RNA-seq) data to define which LFYBS were regulatory, nonregulatory or ‘undefined’. A site was considered to be associated with a significant change in gene expression if it was found in the genomic interval from 3 kb upstream of the TSS to 1 kb downstream of the transcription termination site of a DEG. LFY sites overlapping with genomic regions bound by LFY in ChIP-seq and DAP-seq experiments, and associated with differentially expressed genes, were labeled as ‘regulatory’. The rationale behind this choice is that we wanted to select TFBS that LFY could bind *in vivo* without the requirement for cofactors. LFY sites with no evidence of binding or of differential expression close by were labeled as ‘nonregulatory’. Regulatory and nonregulatory sites were used to train random forest models. All sites satisfying one of the previous criteria but not all of them at once, were named ‘undefined’, as they could not be confidently labeled as either regulatory or nonregulatory.

### Computing genomic features

#### POcc

POcc was calculated using the method published in (Moyroud et al., 2011) and the LFY PWM with dependencies on DNA sequences spanning ± 250 bp or ± 500 bp around each LFYBS. To define the presence of LFYBS around each reference site, we used the same score threshold as to identify LFY sites genome-wide, i.e. the top 0.1% best scoring sites.

#### Computing LFY-other TFs and LFY-LFY distances

To compute the distance between LFYS and other TFBS we used the PWMs of 47 TF clusters from the core plants collection of the JASPAR 2022 database (Castro-Mondragon et al., 2022) to scan the *A. thaliana* genome. We excluded cluster number 47 as it represented a single TF, LFY, which was already included. As we did when defining LFYBS, we computed each cluster’s genome-wide score distribution and set a PWM score threshold at the 99.9th percentile. Then, we used bedtools closest with options -t all -mdb each -d (Quinlan & Hall, 2010) to find the distance, in bp, of each LFYBS from the closest site of each TF cluster. We then transformed such distance to the distance between the central position of each matrix (LFY-cluster) based on matrix length.

To compute LFYBS-LFYBS distances, we used the top 0.1% LFYBS described above and bedtools closest with options -t all -mdb each -d -io (Quinlan & Hall, 2010) to avoid overlaps.

#### Computing LFY-TSS distances

The distance between each LFYBS and the closest TSS was computed using bedtools closest with option -D a (Quinlan & Hall, 2010) to keep positive or negative distance information based on whether the LFY site was upstream or downstream of the closest TSS, respectively. TSS positions were determined based on the TAIR10 annotation (Lamesch et al., 2012).

#### Computing sequence type

We computed six binary features describing the type of sequence where each LFYBS was found: CDS, 5’ UTR, 3’ UTR, intron, promoter and downstream regulatory region. Promoters were defined as 3 kb upstream of the TSS, and downstream regulatory regions as 1 kb downstream of the transcription termination site. Both promoters and downstream regulatory sequences could overlap neighboring genes, thus some genomic regions could be categorized as multiple types at once. For each feature column in the data matrix, ‘1’ indicated that the corresponding TFBS was found in that type of sequence. The TAIR10 gff annotation file was used as a reference (Lamesch et al., 2012).

#### Computing TFBS density and diversity

TFBS density was computed as the total amount of non-LFY TFBS found within ± 250 bp or ± 500 bp around LFYBS.

We used two methods to compute TFBS diversity. The first method was an adaptation of the Shannon entropy formula to quantify non-LFY TFBS diversity as in the following equation:

Equation 1 Shannon’s entropy formula adapted to quantify non-LFY TFBS diversity around LFY sites. *p*_*i*_ is the proportion of non-LFY TFBS of cluster_i_ over the total amount of non-LFY TFBS in the genomic region considered (± 250 bp or ± 500 bp around each LFYBS), and S is the total amount of clusters considered (46 here).

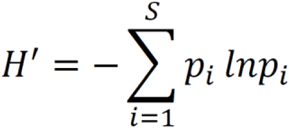

As a second method, we counted the number of different non-LFY TFs with a TFBS within ± 250 bp or ± 500 bp around each LFYBS.

#### Computing average conservation at LFY sites

We used PhastCons and PhyloP conservation scores computed on the *A. thaliana* genome as published in (Tian et al., 2020). Conservation at each position of each LFYBS was retrieved with bwtool extract (Pohl & Beato, 2014) and several strategies were tested to encode PhastCons and PhyloP conservation. The first one was a simple average of the conservation score at each site. The second was a weighted average of the conservation score, where the information content (IC) of each nucleotide in a LFY palindromic PWM (obtained from DAP-seq peaks) was used as weight, depending on its position within the LFYBS. This meant that e.g. if the selected LFYBS sequence had an A in position 1, the IC of the A nucleotide in the LFY’s PWM was used as weight, while if the selected LFYBS had a T in position 1, the IC of the T nucleotide in the PWM was used as weight. The last strategy we used to compute conservation scores was a weighted average using the total IC per position as weight. This meant that a fixed weight was used per position, independently of the nucleotide at each position of the LFYBS sequence.

### Cross-validation and testing scheme

We used the scikit-learn python package to run all machine learning models presented in this paper (v1.2.0) (Pedregosa et al., 2011). First, we split our full dataset following a 80-20 partition. We used 80% of the data as a ‘training set’ to run repeated stratified cross-validation (20 repeats, 5 folds) in order to (i) define the best combination of features to include in the model and (ii) optimize hyperparameters for random forest. The hyperparameters we tested are in Supp. Table 1. Feature combinations are shown in Supp. Table 2 and the best combination is in Table 2.

Once we had defined the best features and best hyperparameters for our random forest, we trained our model on our full training set. We tested its performance on the remaining 20% of our dataset, never seen by the model. Then we used scikit-learn’s function ‘precision_recall_curve’ to plot Precision-Recall curves and their average precision values. The same strategy was used to evaluate random forest model performance after training with specific features.

For logistic regression and SWM models we used the same training-testing split and Precision-Recall curve function. For logistic regression, we used scikitlearn’s LogisticRegression function with the penalty option set to ‘None’, ‘l1’ or ‘l2’. For SVMs, we used scikit-learn’s function SVC with the kernel option set to ‘poly’ or ‘rbf’.

### Feature importance

We used the permutation_importance function implemented in scikit-learn to evaluate the importance of features in our random forest model. As a complementary way to assess feature importance, we also used the property feature_importances_ implemented in scikit-learn random forest models to retrieve Gini importance for each feature in a trained model. We did this during the repeated stratified cross-validation scheme presented in the method section above, on 80% of our dataset.

## Acknowledgments

We thank CNRS for Prime 80 PhD fellowship to L.T. This work was supported by the ANR-17-CE20-0014-01 Ubiflor and ANR-21-CE20-0024 Beflore projects to F.P., the Grenoble Alliance for Integrated Structural & Cell Biology Labex financed within the University Grenoble Alpes graduate school (Ecoles Universitaires de Recherche) CBH-EUR-GS (ANR-17- EURE-0003)

## Author contributions

L.T., A.F., F.P. designed the research, L.T, G.T. performed the plant experiments, F.P., R.B.M., N.T.M, A.F supervised the work, L.T, J.L. performed the bioinformatics analyses, L.T. performed the statistical modeling and wrote the article with the help of all authors.

## Supplementary material

**Supp Fig 1:**
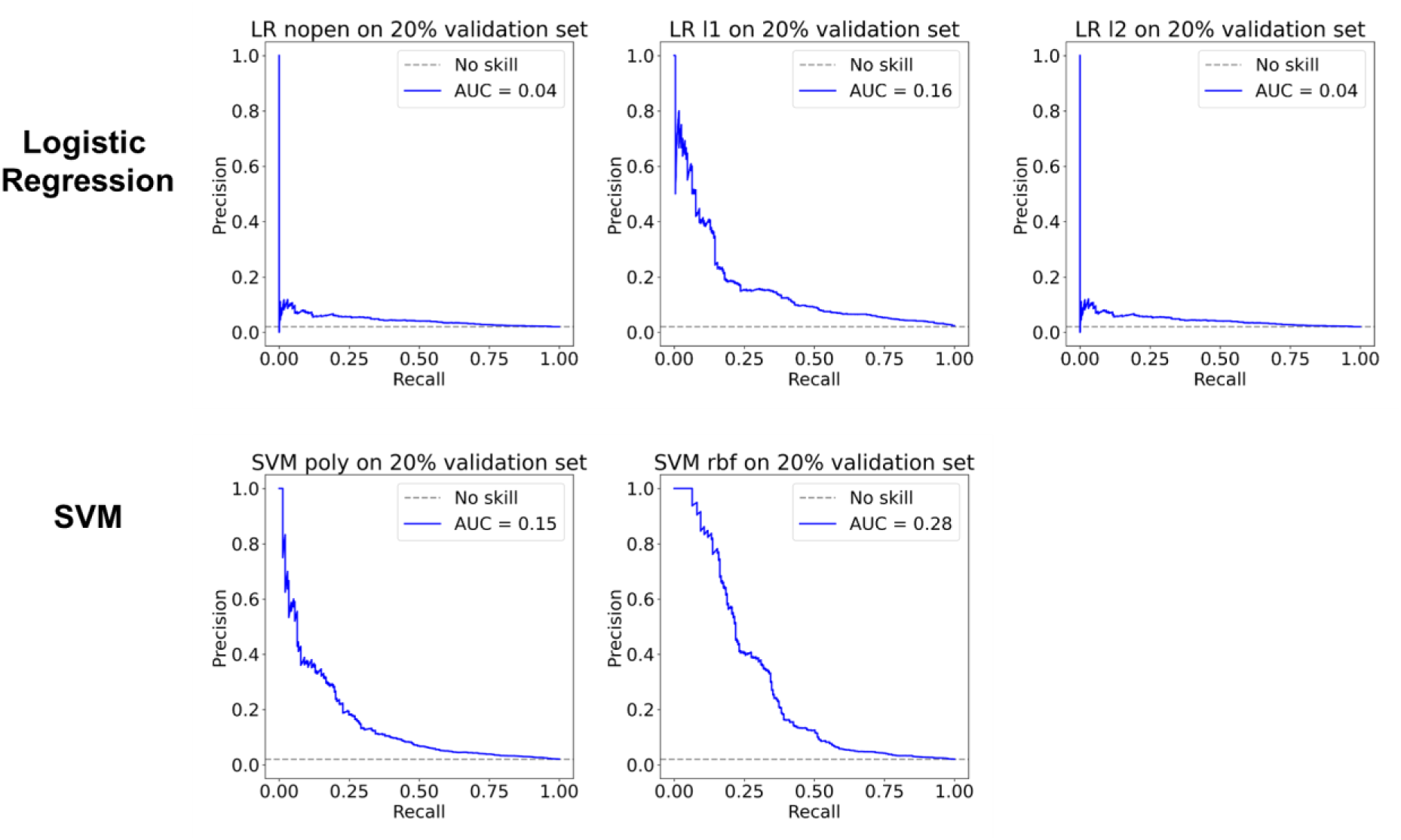
Performance of Logistic Regression and SVM models, evaluated with Precision-Recall curves. Curves were computed on 20% holdout dataset, after training on 80% of our data.

**Supp Fig 2:**
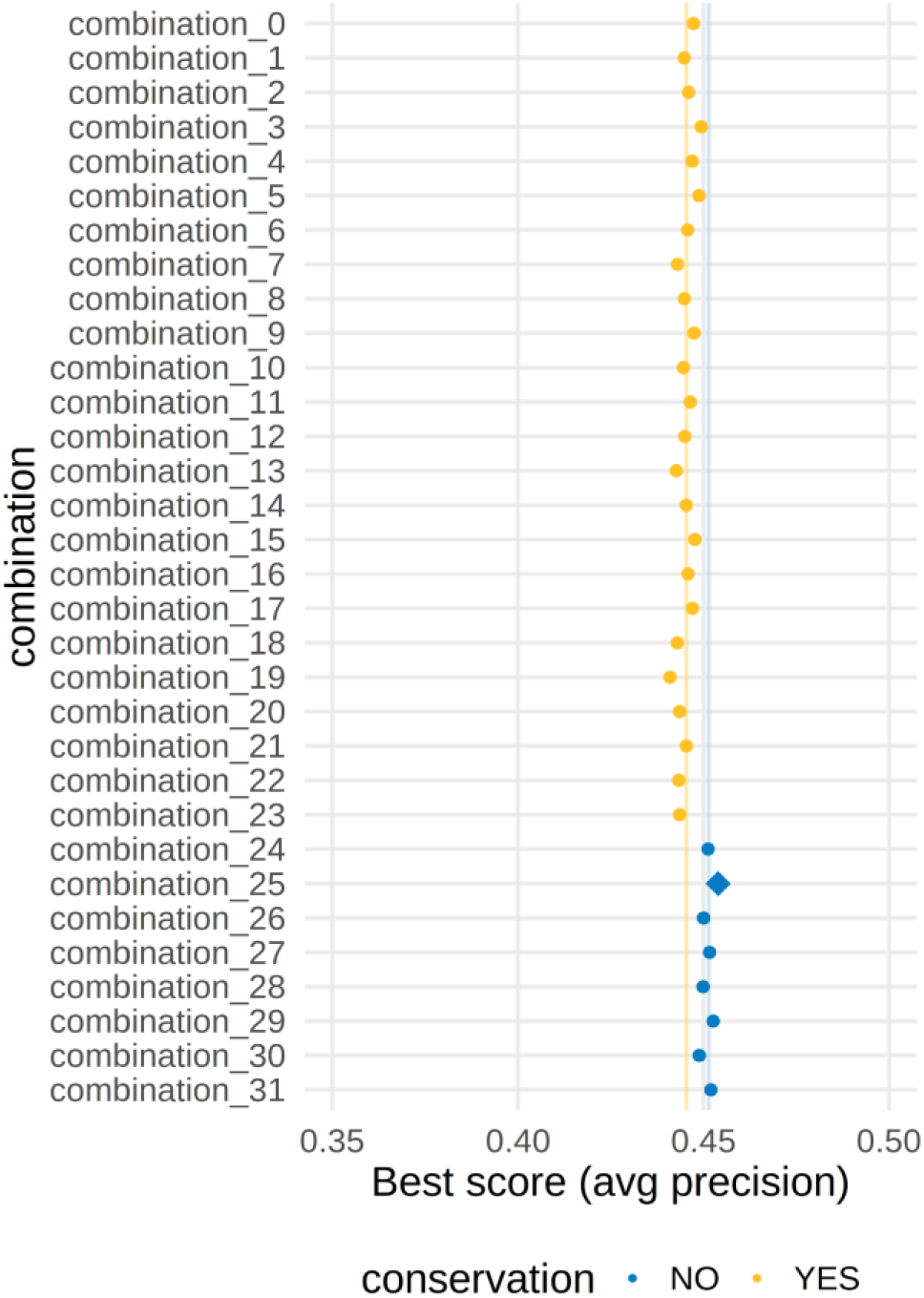
Best average precision score for each combination of features in random forest models obtained through repeated stratified cross-validation and a combination of model hyperparameters. Each dot corresponds to the mean of the average precision score computed with the best combination of hyperparameters for each combination of features. The diamond symbol indicates the combination of features with the highest average precision score. The vertical yellow and blue lines represent the median of scores obtained with feature combinations including and excluding conservation features, respectively.

**Supp. Table 1:** Random forest model hyperparameters we tested. The best combination of hyperparameters is displayed in bold.

| Parameter | Values |
| --- | --- |
| N_estimators | 100, 500, **1000** |
| Max_features | **2**, 7, 10 |
| Criterion | **Gini**, log_loss |
| Class_weight | 'balanced', {0:1, 1:1}, {0:1, 1:2}, **{0:1, 1:10}**, {0:1, 1:20}, {0:1, 1:50} |
| Max_depth | 20, **25**, 30, 35 |

**Supp. Table 2:** Changing features included in each combination. All combinations also included constant features: sequence type, LFYBS quality, LFY-LFY distance, distance of LFY from TFBS of 46 clusters.

| combination | features | best score |
| --- | --- | --- |
| 0 | conservation + Shannon entropy (250) + TFBS quantity (250) + POcc (250) | 0,4473 |
| 1 | conservation* + Shannon entropy (250) + TFBS quantity (250) + POcc (250) | 0,4448 |
| 2 | conservation** + Shannon entropy (250) + TFBS quantity (250) + POcc (250) | 0,4460 |
| 3 | conservation + Shannon entropy (500) + TFBS quantity (500) + POcc (250) | 0,4494 |
| 4 | conservation* + Shannon entropy (500) + TFBS quantity (500) + POcc (250) | 0,4470 |
| 5 | conservation** + Shannon entropy (500) + TFBS quantity (500) + POcc (250) | 0,4488 |
| 6 | conservation + TFBS quantity (250) + total different TFs (250) + POcc (250) | 0,4458 |
| 7 | conservation* + TFBS quantity (250) + total different TFs (250) + POcc (250) | 0,4430 |
| 8 | conservation** + TFBS quantity (250) + total different TFs (250) + POcc (250) | 0,4448 |
| 9 | conservation + TFBS quantity (500) + total different TFs (500) + POcc (250) | 0,4475 |
| 10 | conservation* + TFBS quantity (500) + total different TFs (500) + POcc (250) | 0,4446 |
| 11 | conservation** + TFBS quantity (500) + total different TFs (500) + POcc (250) | 0,4465 |
| 12 | conservation + Shannon entropy (250) + TFBS quantity (250) + POcc (500) | 0,4450 |
| 13 | conservation* + Shannon entropy (250) + TFBS quantity (250) + POcc (500) | 0,4427 |
| 14 | conservation** + Shannon entropy (250) + TFBS quantity (250) + POcc (500) | 0,4454 |
| 15 | conservation + Shannon entropy (500) + TFBS quantity (500) + POcc (500) | 0,4477 |
| 16 | conservation* + Shannon entropy (500) + TFBS quantity (500) + POcc (500) | 0,4459 |
| 17 | conservation** + Shannon entropy (500) + TFBS quantity (500) + POcc (500) | 0,4471 |
| 18 | conservation + TFBS quantity (250) + total different TFs (250) + POcc (500) | 0,4429 |
| 19 | conservation* + TFBS quantity (250) + total different TFs (250) + POcc (500) | 0,4410 |
| 20 | conservation** + TFBS quantity (250) + total different TFs (250) + POcc (500) | 0,4436 |
| 21 | conservation + TFBS quantity (500) + total different TFs (500) + POcc (500) | 0,4454 |
| 22 | conservation* + TFBS quantity (500) + total different TFs (500) + POcc (500) | 0,4433 |
| 23 | conservation** + TFBS quantity (500) + total different TFs (500) + POcc (500) | 0,4436 |
| 24 | Shannon entropy (250) + TFBS quantity (250) + POcc (250) | 0,4512 |
| 25 | Shannon entropy (500) + TFBS quantity (500) + POcc (250) | 0,4539 |
| 26 | TFBS quantity (250) + total different TFs (250) + POcc (250) | 0,4500 |
| 27 | TFBS quantity (500) + total different TFs (500) + POcc (250) | 0,4516 |
| 28 | Shannon entropy (250) + TFBS quantity (250) + POcc (500) | 0,4499 |
| 29 | Shannon entropy (500) + TFBS quantity (500) + POcc (500) | 0,4526 |
| 30 | TFBS quantity (250) + total different TFs (250) + POcc (500) | 0,4489 |
| 31 | TFBS quantity (500) + total different TFs (500) + POcc (500) | 0,4520 |

**Supp Fig 3:**
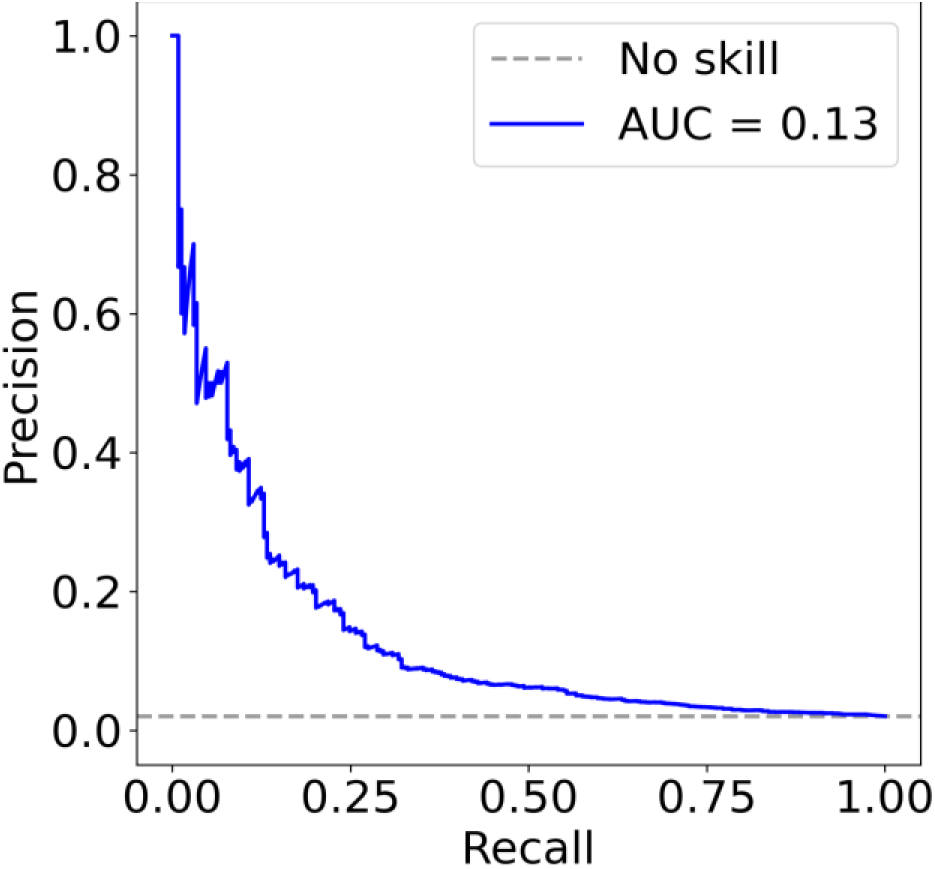
Precision-Recall curve obtained with a model trained exclusively with three most important features (POcc, PWM and LFYBS-LFYBS). Training was performed on 80% of data, testing to compute this curve was performed on the remaining 20%. Dashed gray line represents the performance of a random model.

**Supp Fig 4:**
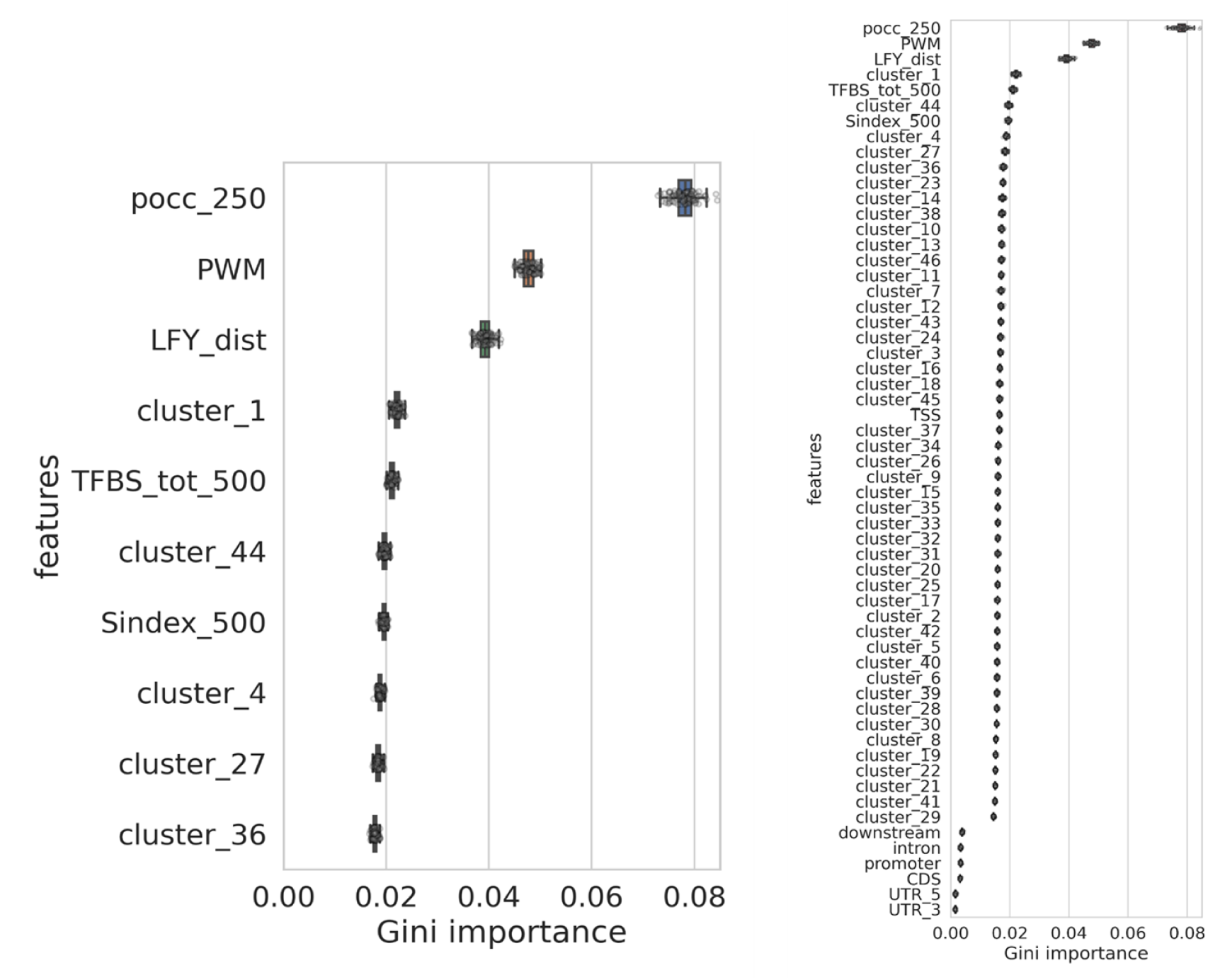
Gini importance as a measure of feature importance. Top ten features are displayed on the left, all features on the right.

**Supp Fig 5:**
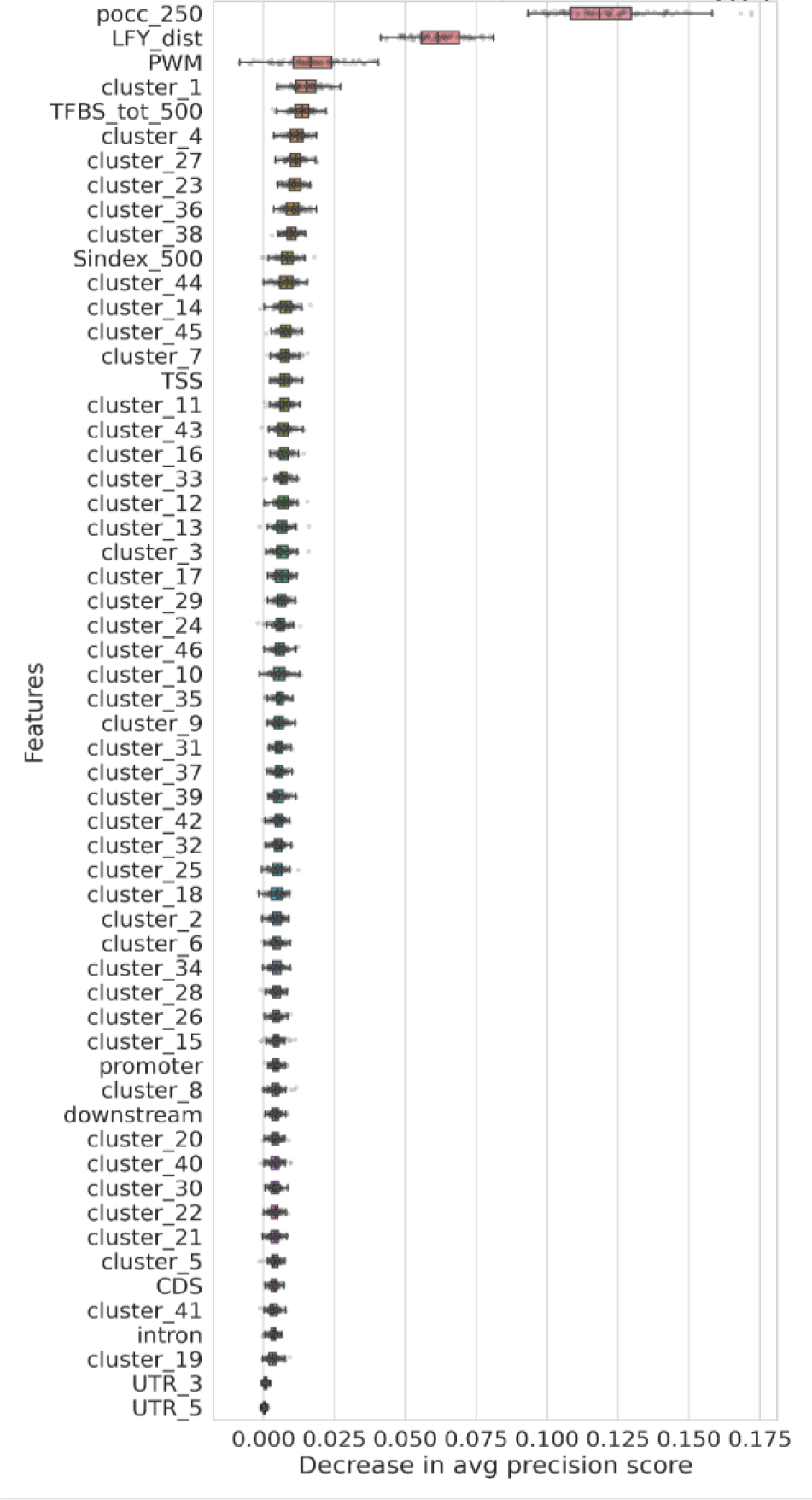
Permutation importance, expressed as the decrease in average precision score, as a measure of feature importance. All features included in the model are shown.

## Notes

### Competing Interest Statement

The authors have declared no competing interest.

